# A theoretical high-density nanoscopy study leads to the design of UNLOC, an unsupervised algorithm

**DOI:** 10.1101/275313

**Authors:** Sébastien Mailfert, Jérôme Touvier, Lamia Benyoussef, Roxane Fabre, Asma Rabaoui, Marie-Claire Blache, Yannick Hamon, Sophie Brustlein, Serge Monneret, Didier Marguet, Nicolas Bertaux

**Author notes:** Correspondence should be addressed to DM or NB. These authors jointly supervised this work (Didier Marguet, Nicolas Bertaux).

## Abstract

Among the superresolution microscopy techniques, the ones based on serially imaging sparse fluorescent particles enable the reconstruction of high-resolution images by localizing single molecules. Although challenging, single-molecule localization microscopy (SMLM) methods aim at listing the position of individual molecules leading a proper quantification of the stoichiometry and spatial organization of molecular actors. However, reaching the precision requested to localize accurately single molecules is mainly constrained by the signal-to-noise ratio (SNR) but also the density (*D_frame_*), i.e., the number of fluorescent particles per μm^2^ per frame. Of central interest, we establish here a comprehensive theoretical study relying on both SNR and *D_frame_* to delineate the achievable limits for accurate SMLM observations. We demonstrate that, for low-density hypothesis (i.e. one-Gaussian fitting hypothesis), any fluorescent particle biases the localization of a particle of interest when they are distant by less than ≈ 600 nm. Unexpectedly, we also report that even dim fluorescent particles should be taken into account to ascertain unbiased localization of any surrounding particles. Therefore, increased *D_frame_* quickly deteriorates the localization precision, the image reconstruction and more generally the quantification accuracy. The first outcome is a standardized density-SNR space diagram to determine the achievable SMLM resolution expected with experimental data. Additionally, this study leads to the identification of the essential requirements for implementing UNLOC (UNsupervised particle LOCalization), an unsupervised and fast computing algorithm approaching the Cramér-Rao bound for particles at high-density per frame and without any prior on their intensity. UNLOC is available as an ImageJ plugin.

---

Cellular processes rely on the proper stoichiometry and coordinated spatial organization of molecular actors. In this regard, single-molecule localization microscopy (SMLM) provides reliable information on molecular distributions, interactions and biological processes by counting and localizing single fluorescent particles (1, 2). As such, by aiming at listing the position of single molecules (3-5), SMLM differs significantly from the superresolution microscopy techniques which generate reconstructed images based on density distribution of the fluorescent signals (6, 7).

SMLM imaging is of low technical complexity but ensuring the robustness of the super-resolution observations requires a rather high attention for rigorous sample preparation, image recording and data analysis (2, 8). In a first attempt, SMLM imaging is best designed at describing the molecular organization in fixed biological samples to overcome impaired localization precision due to molecular diffusion during the camera exposure for any dynamic processes occurring in living cells (9). To a lesser extent, insufficient sample fixation alters for the same reason the localization precision and consequently, the final resolution of the reconstructed images (8, 10). SMLM imaging also requires the design of well-defined fluorescent labeling probes to preserve the localization precision (9, 11). This implies to ascertain the probe specificity, to minimize the steric constraint or to optimize the dark/fluorescent state ratio (12). Finally, the quality of SMLM imaging strongly relates to a robust data processing.

Based on the theoretical principle governing the accurate localization (13-16), SMLM analyses were initially performed with algorithms fitting the point spread function (PSF) of imaged isolated particles (17). In essence, accurate localization with a one-Gaussian fitting hypothesis requires effective low-density (LD) imaging conditions (i.e., low *D_frame_*;Fig. 1a, left panel). However, such conditions intrinsically undergo frequent experimental exceptions. Possible high local densities, even at a low density per frame, are encountered due to a non-homogeneous distribution of the particles or to the stochastic nature of SMLM methods. Moreover, complex variations of the background are very frequently observed both in space and time. Altogether, this fully justifies to quantify SMLM data with an accurate HD mode of analysis.

A canonical scenario with two particles recapitulates the problem (Fig. 1a, middle panel). Postulating LD conditions on these data generates not only a lack of precision but also a bias regarding their localizations. The retrieval of accurate precision and unbiased localization data can be obtained by a two-Gaussian fit under a high-density (HD) hypothesis. When the two particles are too close to each other (Fig. 1a, right panel), the scenario becomes non-resolvable (NR). The HD hypothesis is no longer a working scenario, as shown by indistinguishable fitting residues (Fig. 1b). Thus, an accurate enumeration and localization of particles rely not only on the signal-to-noise ratio (SNR) but also on the density of particles per frame (*D_frame_*). Other analytical approaches have been developed for HD SMLM data such as the ones based on compressed sensing (18, 19), Bayesian estimation (20, 21) or deconvolution (22) analyses. They provide the local density of the emitting particles, but without reaching the single-molecule localization precision obtained by the multi-Gaussian fitting methods (23-25) (see also (26) for review).

Assessing accuracy and robustness for localization precision requests first to define theoretically the reliability of SMLM observations. By simulating realistic conditions, we investigate a theoretical canonical scenario with two particles on the localization precision. We characterize the bias on the localization precision based on an appropriate estimator and the Cramér-Rao bound (CRB). We demonstrate the impact of the SNR and *D_frame_*on the localization precision. Our theoretical study prompted us to develop original tools to estimate and ensure accurate quantitative SMLM analyses:(i) a unique density-SNR space diagram that enables standardized evaluation of the localization accuracy expected from experimental data and (ii) UNLOC, a real parameter-free, unsupervised and fast computing algorithm as a plugin for ImageJ (27) especially useful for any inexperienced researchers requiring a rigorous SMLM quantification.

**Figure 1.**
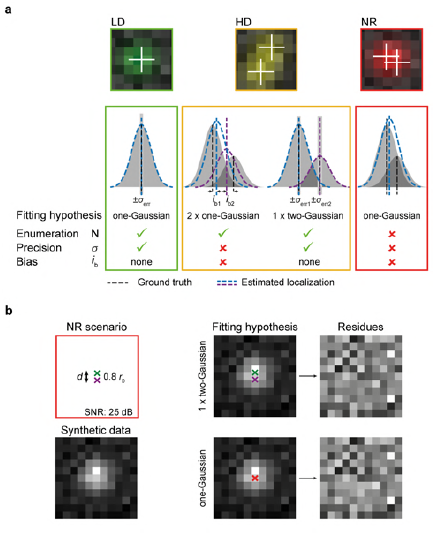
│ Enumeration, precision and bias for single-molecule imaging. (**a**) Canonical scenarios for one (effective LD conditions) or two particles (HD and NR conditions) with the corresponding fits obtained using a one- or two-Gaussian hypothesis and the incidence of the enumeration and localization precision ± bias. (**b**) NR scenario analyzed using an LRT (PD ≈ 0 at PFA = 10^−4^ and PD = 0.2 at PFA = 10^−2^), with indiscernible differences in the residues between the two hypotheses (see Note S3).

## MATERIALS AND METHODS

### THEORETICAL FRAMEWORK CONSIDERATIONS

The theoretical statistical study is based on a general signal-dependent noise model of the intensity *x*_*p*_ at pixel *p* = (*i, j*) as a proper model matching the stochastic processes that occur during experimental acquisition of SMLM (Note S1, Eq. S1). Also, for the sake of generalization, we express the inter-particle distance *d* by the characteristic dimension r_0_ of the PSF and the SNR in a logarithmic decibel scale (SNR_dB_ = 10 _log10_ SNR). All derivations and analyses are detailed in Notes S1-5 and Table S1 for acronyms and symbols used in this study.

#### PSF size

We hypothesize that the PSF exhibits a classical integrated Gaussian profile with a characteristic size *r*_0_ represented by the full width at half maximum (FWHM) of the peak according to:

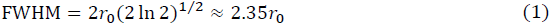

For a comprehensive order of magnitude, the *r*_0_ value is equal to 130 nm for an optical pixel size of 107 nm (r_0_ = 1.25 *pixels*). For this characteristic size, 86% of a single signal power is within 3 × 3 pixels, with the power defined as Σ_*p*_(αg_p_)^2^ with α, the signal intensity at the pixel *p*

#### Definition of a single contrast parameter

A pertinent contrast parameter should describe the difficulty of the task, i.e., the expected precision, independently of the data to ascertain the same precision of particle localization for data acquired on different equipment. We chose the SNR as an efficient contrast parameter, which was calculated using the following equation assuming that ∬ *g*^2^(*x, y*)*dx dy*= 1:

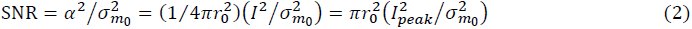

where *α* is the amplitude of the signal, 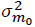 the variance of the fluorescent background *m*_0_, *I* the intensity of a particle (in number of counts), *I_peak_* the peak signal intensity, and *r*_0_ expressed in pixel units. As compared to the classical contrast parameters defined by the sole signal intensity or by the ratio of the particle signal to the background level, the SNR alone recapitulates the expected achievable precision (Fig. S1). The performance limits are fixed by the CRB, which, for a given density D _*frame*,_ depends on (SNR, *m*_0_, *G*), but is only recapitulated by the SNR:

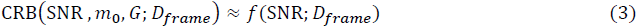

with 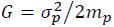 the electron-multiplying (EM) gain which differs from the electron-multiplying charge-coupled device (EMCCD) gain sets experimentally.

#### Signal-dependent noise model

An EMCCD or complementary metal-oxide semiconductor (CMOS) camera is designed for low-light imaging. However, regular experimental SMLM image acquisition deviates significantly from these conditions and covers a wide range of intensities, from a few hundred to thousands of photons in the background and signal areas (Note S1). Since the PSF corresponds to the averaged repartition of photons for spatially isolated punctate signals, the particle intensity contributes to the SNR and the quality shape of the detected peak. A precise particle localization implies that an appropriate SNR has been achieved and, hence, a sufficient photon flux recorded during the integration time.

We consider a general signal-dependent noise model of the intensity *x*_*p*_ at pixel *p*= (*i,j*) as a realistic model of the signal fluctuations that approximates these conditions:

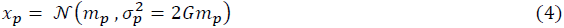

where *m*_*p*_ is the averaged intensity for pixel *p*, and *σ*_*p*_ is its standard deviation. This equation has been corroborated by others (28).

#### Choice of the MMSE as appropriate estimator

Compared to a maximum likelihood estimator (MLE), the use of a minimum mean square error (MMSE) estimator has received increasing interest (17, 29, 30). Under LD conditions, the MMSE estimator corresponds to a simple filtering operation (Note S2). Under HD conditions, the two estimators show a similar mean square error (MSE) that is similar to the CRB, as well as a very limited bias in cases in which two particles P_1_ and P_2_ are present at variable distances or at variable SNRs but at a constant SNR difference (Δ_SNR_) (Fig. S2). However, the MLE is more biased than the MMSE estimator when they are compared under a defective LD hypothesis or for images with a non-homogeneous background (Figs. S3a, and S8, Note S5). Moreover, an MMSE estimator remains less complex than an MLE, which requires optimizing Eq. S25 and, consequently, a higher computational cost.

### GENERATION OF SYNTHETIC DATA

All synthetic data were generated with Matlab (The MathWorks Inc., Natick, MA). Images must have met the conditions of a signal-dependent noise model, as defined in Eq. 4, to achieve a controlled scenario. The PSF of unitary power is defined by three parameters, *SNR, r*_0_ and *w*_*n*_, where is the size of the window support. When it is requested, *d* or *D_frame_* were set. Otherwise specified, the synthetic data were generated based on the following values: *r*_0_ = 1.25 pixels, *M*_0_ = 300, *G* = 1, a pixel size of 107 nm and *SNR* _*dB*_ = 25 *dB*.

## UNLOC ALGORITHM

### Description

UNLOC provides a list of coordinates and associated parameters for each detected particle for *a posteriori* quantification and image reconstruction. The algorithm is based on the decision theory, which only requires the PFA value to be set without the initialization of any parameters relative to the data (SNR, particle density, or background level). UNLOC is an iterative algorithm that alternates an overestimation of the number of particles to obtain a minimal fitting residue with a general likelihood ratio test (GLRT) to suppress useless particles. UNLOC calculates the position errors based on the CRB for particles at high density per frame and without any prior on their intensity. The UNLOC principle is illustrated in Figs. 4 and S9 and the mathematical details are provided in Note S6.

#### Performance evaluation

We evaluate UNLOC performances on (i) localization accuracy, (ii) inhomogeneous density distribution and (iii) computation time.

##### Localization accuracy on synthetic data

A first chart assesses the limits for a simple case with two particles at different SNRs (from 20 to 30 dB) and variable inter-particle distances (from 0.5 to 6 *r*_0_) (Fig. S11); the alphanumeric characters and *r*_0_ scales were generated under LD conditions to avoid any interference with the evaluated signals. The second chart reproduces the density-SNR space diagram with the alphanumeric character defined by an SNR and a local density ranging from 20 to 40 dB and 0.1 to 5 part/μm^²^/frame, respectively. Briefly, each character is designed by dots in a 6 × 12 subpixel area with a subpixel size of 0.1 pixel. The local density is mimicked by the illumination rate of subpixels: the more dots of a given character are illuminated simultaneously, the higher is the local density.

**Figure 4.**
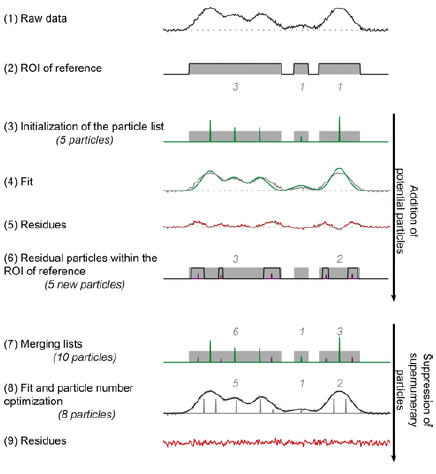
│ Principle of UNLOC. Global heuristic of the UNLOC HD mode used for detection and enumeration. Regions of interest in the image containing potential particles are defined through a GLRT and a first list of particles is created (steps 2-3). A first multi-Gaussian fit is performed (step 4) and a new list of particles is created based on the residues (steps 5-6). Following the addition of potential particle steps, supernumerary particles are suppressed by an optimization loop of multi-Gaussian fits, GLRT on residues to find the number of particles and their respective positions, sizes and intensities (steps 7-9).

##### Inhomogeneous density distribution

Synthetic data mimicking clustered molecules by varying cluster and particle densities (3 to 10 clusters/μm^²^ and 0.1 to 1 part/μm^²^/frame) are generated. Clusters of ≈ 97 particles with a 100 nm diameter at 27 dB of SNR were randomly dispersed in a 10 × 10 μm image (i.e., 94 × 94 pixels), resulting in a variable total number of particles. The UNLOC results and ground truth data were analyzed using two clustering algorithms, a standard clustering algorithm, DBSCAN (31), and a dedicated algorithm for SMLM data, SR-Tesseler (32) (Table S2).

##### Computation time

This relates to the hardware characteristics and data complexity in particle density/frame, SNR, number of pixels and frames. All synthetic and experimental data were computed on a Dell Precision T7910 with a 2 × 12 cores Intel^®^ Xeon^®^ processor (E5-2687W v4, 3 GHz) and 128 GB of RAM.

#### Software package

The UNLOC ImageJ plugin, including a user’s manual and the scripts for the charts is freely available for academic and nonprofit use as supporting software package or at http://ciml-e12.univ-mrs.fr/App.Net/mtt/ (including updated versions).

## SAMPLE PREPARATION, MICROSCOPY SETUP AND DATA ACQUISITION

dSTORM images of Alexa Fluor^™^ 647-phalloidin (Thermo Fisher Scientific)-labeled actin filaments in fixed COS-7 cells (ATCC CRL 1651) were prepared using the method reported by Xu et al. (33). For imaging, samples were mounted on a single-depression concave slide (VWR # 630-1611) in dSTORM buffer (50 mM Tris-HCl, pH 8.5, 50 mM cysteamine and 50 mM NaCl in 18.2 MΩ.cm Milli-Q water) and sealed with Picodent Twinsil speed 22 (Picodent, Germany).

Acquisitions were performed on a custom-built microscope with an excitation light path for total internal reflection fluorescence (TIRF) and wide field observations (Fig. S14 and Table S3). Samples were illuminated with a 647 nm laser at 4 kW.cm^-2^. Typically, ≈ 57,000 frames were acquired at an exposure time of 36 ms and an EMCCD gain of 100. The axial drift was corrected by the autofocus module. Data were analyzed by UNLOC in HD mode with a high spatial frequency variation of background, a reconnection process with one Off-state lifetime frame and an integrated Gaussian rendering process after drift correction by correlation and without data filtering.

## RESULTS AND DISCUSSION

### THEORETICAL STATISTICAL STUDY

The SMLM analytical methods can be divided into two subgroups. A first one is primarily used for low-density data (13-16, 34) and intends to nearly achieve the CRB performances using a MLE (29, 35). A second subgroup is specifically designed for high-density data set or fast computing; it reconstructs images without necessarily achieving the best detection performance or localization precision (18-22). Combining the benefits of these two subgroups should maintain fast processing for unbiased particle localizations by approaching the CRB under various SNR and *D_frame_* conditions.

We define the SNR as a pertinent contrast parameter (Fig. S1 and Note S1) and *D_frame_* as the number of fluorescent particles per μm^²^ per frame. *D_frame_* is linked to the inter-particle distance distribution (36) that is implicitly related to the probability *p* of finding a single particle at a radius *r* for randomly distributed particles, as determined using the following equation:

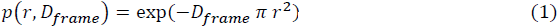

A canonical scenario with two particles recapitulates the problem by identifying the intrinsic limits of the LD/HD and HD/NR transitions (Fig. 1) to design a heuristic adapted to divers experimental data.

Here, we present a theoretical study using two particles, P_1_ and P_2_, of respective intensities *α*_1_ and *α*_2_, which are separated from each other by *d*, the inter-particle distance expressed in *r*_0_, the characteristic PSF dimension.

#### Limit of the LD hypothesis

According to LD hypothesis, the localization precision is governed by a photon-counting analysis, regardless of the local particle density (13). Therefore, the omission of surrounding particles introduces a bias, *i*_*b*_, into the localization of particle P_1_. By considering a signal-dependent noise model (28) (Note S1),*i*_*b*_ may be solved for an MMSE estimator within an analysis square window of dimension *w* as follows (Note S2):

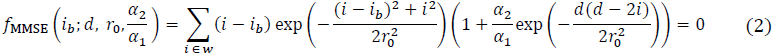

The LD/HD transition distance at which the bias becomes greater than the expected precision of the localization of P_1_ also relies on the intensity ratio between the two particles (Fig. 2a). This result is validated using synthetic data, where the P_1_ position is estimated with an MMSE estimator initialized at its true coordinates (Figs. 2c, S3). We set *d*_*LD*/*HD*_, the LD/HD transition distance, at a large dynamic intensity ratio ((*α*_2_/*α*_1_)dB = 22 dB), a scenario that is conceivable based on experimental data (Fig. 2b). This transition occurs at approximately 5*r*_0_ [i.e., ≈ 650 nm for a standard *r* _0_ ≈ 130 nm]. Thus, any particles spaced at a distance below this limit require an HD estimation.

**Figure 2.**
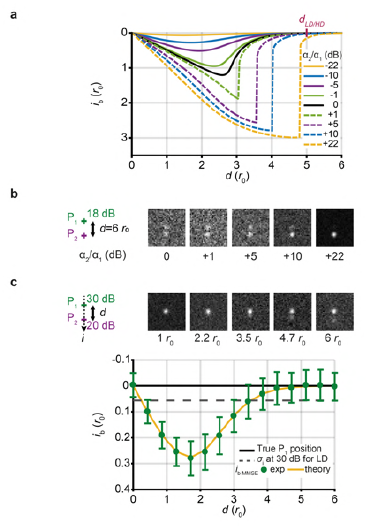
│ Theoretical limit of the LD hypothesis and its validation on synthetic data. (**a**) Theoretical bias for P1 localization using an MMSE estimator under the LD hypothesis at different intensity ratios, *α*_2_/*α*_1_ (*w*= 9 pixels, *r*0 = 1.25 pixels [≈ 130 nm]). (**b**) Synthetic data for two particles at different intensity ratios, *α*_2_/*α*_1_.(**c**) Bias *i*_*b*_ for P_1_ using an MMSE estimator on synthetic data (see also Fig. S1). Mean ± s.d., n = 250 images per *d* value.

#### Limit of the HD hypothesis

Now, we must primarily consider HD conditions to determine the HD/NR transition, i.e., the distance from which an enumeration is no longer achievable (Note S3). We determine this limit by performing a likelihood ratio test (LRT) that yields the best probability of detection (PD) set at a probability of false alarm (PFA) (37). For a simple NR scenario (Fig. 1b), two particles separated by a short distance have an almost null PD at the usual PFA (10^−4^). Similar to LD cases, the enumeration error biases the localization of the other particles due to the erroneous fitting hypothesis. This detection error alters the localization of the nearest and more distant particles (Fig. S4a). Conversely, the detection of any dim particles enables unbiased localizations (Fig. S4b). Of note, both SNR and *D_frame_* contribute to the assignment of a proper PD. Hence, the HD/NR transition *d*_*HD /NR*_ (SNR) must be set to given SNR and appropriate PD, e.g., *d*_*HD /NR*_ (25 dB) = 1.23 *r*_0_[≈ 170 nm] at a weakly restrictive PFA (10^−3^) for 80% PD (Note S3, Fig. S6). Moreover, the nuisance effect propagates at long distances and depends on both the SNR and the geometry. Thus, minimization of the bias in the localization of bright particles implies that even the particles with a weak intensity must be taken into account in the initial detection step.

#### LD, HD and NR subsets

Based on Eq. 1, which provides the probability of identifying isolated particles in a random distribution and at the inter-particle distances delineating the LD/HD and HD/NR transitions, we easily obtain the partition of the LD, HD and NR subsets as follow:

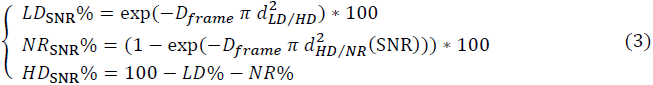

Significantly, at a SNR of 25 dB, the NR subset rapidly increases with *D_frame_*, to ≈ 10% at 1 part/μm^²^/frame and up to ≈ 40% at 5 part/μm²/frame (Fig. 3a). This subset is quantifiable for a known scenario (simulated data) but is obviously not quantifiable for experimental data, leading to significant erroneous particle quantifications. Thus, we determine the densities/frame corresponding to the respective thresholds *d*_*LD/HD*_ and *d*_*HD/NR*_ (SNR) at a suitable quantile Q_*subset*_ set to 20%, which refers to the densities at which 20% of the particles have a *d* value below these thresholds (Fig. S5a).

**Figure 3.**
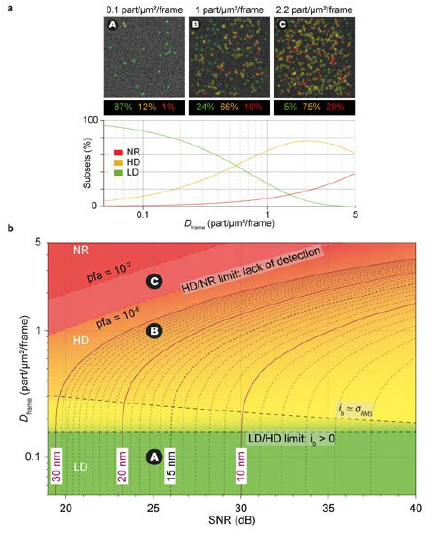
│ Theoretical and SNR limits. (**a**) Color-coded LD, HD and NR subsets as a function of with ground truth thumbnails at different densities (SNR = 25 dB, *r*_0_ ≈ 130 nm and 10^−4^ PFA for the HD/NR transition). (**b**) Density-SNR space diagram for data at a known *r*_0_ combining the transition limits (color-coded) and precision accuracy (isocurves at Q_σ_ = 80%). The thumbnails in (**a**) are located on the diagram. A: 87% of particles were localized with a σ_RMS_ ≤ 17 nm. B: 80% of particles were localized with a σ_RMS_ ≤ 26 nm. C: 20% of particles are non-resolvable at 10^−3^ PFA.

#### Merged information in a single density-SNR space diagram

We generalize this canonical two-particle study to the achievable limits for accurate enumeration and localization in realistic *D_frame_* and SNR ranges, and in the case of local homogeneous particle distribution. If the SNR predominantly determines the expected localization precision σ_*RMS*_ under LD conditions, this postulate is no longer valid under HD conditions where increased *D_frame_* decreases σ_*RMS*_ (Fig. S5b). From local density, we identify three limits (Fig. S6, Note S3):

(i) *a precision limit* when the error becomes greater than the expected localization precision. We would like to point out that calculating the CRB assuming known the signal intensity or at least a prior on its value has no impact on the localization precision for LD data. However, for HD data, the CRB have been calculated in previous studies without estimating the particle intensity (14, 38, 39), leading to an overestimation of the actual resolvable *D_frame_*. In fact, we demonstrate that only a very precise knowledge of the particle intensity improves the localization precision but such requirement is actually unrealistic for experimental data (Note S4). Therefore, we calculate the CRB by jointly evaluating the position and intensity of the particles;

(ii) *a separation limit* when the errors of the particle positions become larger than the distance separating them;

(iii) *a detection limit* when, at a given PFA and reasonable PD, the number of particles can no longer be determined.

Although these limits have the same order of magnitude at same SNR (Fig. S6), the detection limit is the effective limiting factor since, by no longer achieving accurate particle detection, the scene becomes non-resolvable. Therefore, we define the inter-particle distance limit *d*_*HD/NR*_ (SNR) for HD/NR transitions as the distance at which an optimal detector with known values for all parameters (positions, intensity, noise and background) has a PD of 80% at PFA = 10^−3^ (Figs. S5a, S6b). Moreover, based on the cumulative σ_*RMS*_ distribution (Fig. S5b), we arbitrarily set a quantile *Q_σ_* to 80% to plot the precision isocurves: for a given *D_frame_*, 80% of particles have aσ_*RMS*_ less or equal to the precision isocurve value.

Finally, we merge the color-coded information for LD/HD/NR transition areas with the precision isocurves in a density-SNR space diagram (Fig. 3b). This diagram defines the experimental local *D_frame_* and SNR required to reach a given precision. For example, a 15 nm precision is achievable at 26 dB under effective LD conditions (*D_frame_* <0.15 part/μm^²^/frame). At 1 part/μm^²^/frame, this precision requires data obtained at 30 dB. At a two-fold higher *D_frame_*, only the particles with high SNR (≥ 40 dB) are localized with this precision. The SMLM performance also decreases when *r*0 must be estimated (e.g., wide field versus TIRF microscopies, Fig. S7).

## UNLOC – UNsupervised particle LOCalization

### UNLOC Principle

Our theoretical study allowed us to conceive UNLOC, an algorithm based on an unsupervised heuristic, while minimizing the computational cost and improving the analytical reliability of complex data (Note S6). Additionally, UNLOC is designed to achieve the best performances, as delineated here by the CRB, for any particles separated by a distance greater than *d*_*HD/NR*_ (SNR) and without any prior on their intensity.

Although UNLOC accounts for spatiotemporal variations in the background, SNR and local particle density, it only requires the PSF size (*r*_0_) of the microscope to build a list of particles. The algorithm is divided into three modules (Fig. S9a): (i) a detection/estimation module performed in an iterative process (Fig. 4) together with optimization of the number, position and intensity of the particles, and also the PSF size if requested (Fig. S9b); (ii) a reconnection module to minimize an overrepresentation of particles in the reconstructed image using statistical tracking of the particle history over the frames;(iii) an optional module for drift correction and classical image rendering methods.

#### Detection/estimation

Two modes of detection/estimation are provided. A standard mode appropriate under effective LD conditions is similar to MTT (40). It is based on a GLRT at unknown background which is an efficient unsupervised detector that only depends on the PFA (41). Alternatively, the HD mode of UNLOC should be applied for any data with high local density/frame and complex background, a condition which occurs very frequently with experimental data. Preserving the GLRT robustness under HD conditions requires an estimation of the background mean and variance. This is done by filtering and interpolating the background between the regions of the frame without signals.

The HD module establishes a list of particle localizations by the iterative addition/suppression of the number of particles (Figs. 4, S9b). It initially provides a robust estimate of the mean and variance of the background in a single frame or within a set of consecutive frames when a high density of particles and important variations are observed in the background between frames. Thus, the initial regions of interest (ROI_ref_) comprising signals are detected with a GLRT at known background parameters and at a PFA of 10^−6^, a threshold which ascertains almost a PD of 100% for dim particle (SNR ≥ 18 dB) (37, 41, 42). A list of particles is initially generated by assessing the maxima in each ROIref on the deconvoluted frame. ROI_ref_ are classified into two types of subgroup (43): one encompasses all isolated particles that only requires localization using a one-Gaussian fit, and the other encompasses all ROI_ref_ in which at least two particles are present at *d* <5*r*_0_. Since any dim particle can introduce a localization bias, these subgroups must contain any detected particles, regardless of their intensity. Each subgroup is further analyzed using an iterative process to identify new ROIs present within the residual fit of the initial ROI_ref_. The list of additional particles identified according to their maxima is merged with the previous list. Next, a global fitting procedure performed using a GLRT suppresses supernumerary particles from the list. The enumeration procedure is executed conjointly with the optimization of particle position, intensity and PSF size.

#### Reconnection

If the detection/estimation step is designed for both static (SMLM on fixed samples) and dynamic (single-particle tracking (SPT)) observations, the reconnection step applies solely to static observations within the intrinsic theoretical limits (see (40) for SPT dedicated algorithm). To avoid an overestimation of the molecule count, fluorescent signal over consecutive frames or over disconnected frames due to the reversible switch between the fluorescent and dark states should be reconnected (Note S6, paragraph SN 6.3). We define the statistical tracking of the particle history over the frames as a “trajectory”. Thus, live trajectories (i.e., trajectories identified in a restricted number of previous frames) are tested using the particles present in the current frame. We define *S*_PD_ as a threshold set by the expected probability for the reconnection domain and the variances of both the trajectory precision and the nearest particle. Moreover, a reverse test (particle to trajectory) validates the reconnection.

#### Drift correction, data filtering and rendering

Drift correction is achieved by two optional methods: (i) automated tracking of fluorescent fiduciary markers (3) over the frames using a GLRT or (ii) image cross-correlation (44) in which subsequent resolved frames are correlated. Standard data filtering is implemented to remove outliers (e.g., any data with low precision in localization, intensity, SNR, ON-state, etc.). Finally, five classical rendering modes (45) are implemented to reconstruct the SMLM image: (i) a binary mode that represents each localization as a white sub-pixel in the reconstructed image, (ii) an integrated binary mode in which the intensity values assigned to each sub-pixel correspond to the number of localizations within each pixel, (iii) a time mode encoded in a look-up table as a function of the first temporal appearance of the particle, (iv) a mode where each particle is represented by a Gaussian whose the variance reflects the localization precision and (v) an integrated Gaussian mode similar to the previous mode but accounting for the local density.

### UNLOC performances

Although the super-resolution algorithm evaluation mostly considers the final image resolution (30, 46, 47), we favor at benchmarking the localization precision as the primary goal ensuring accurate SMLM data. Classically, benchmarking metrics pair the ground truth of synthetic bio-inspired data and the estimated particle localizations at a particular tolerance radius (26). If these metrics are pertinent under real LD conditions, they have unsolvable solutions at increased *D_frame_* (Fig. S10), with mismatches resulting from NR particles or inappropriate algorithm outputs. Usually, the tolerance radius is the same as the PSF FWHM value (26) and does not account for the localization precision. At a homogeneous *D_frame_* of 2 part/μm^²^/frame, 50% of the particles are located at a closer distance than this tolerance radius, providing inconsistent pairings. For this reason, the UNLOC performances are directly visualized on two charts (Fig. 5): one for assessing the limits for canonic two-particle scene and the other for mimicking the density-SNR space diagram. The UNLOC analysis of the inter-particle distance chart explicitly clearly verifies the main conclusions of our theoretical study. In LD mode, a bias starts occurring for any particles distant from less than around 5 *r*0, i.e.,*d*_*LD/HD*_. Another bias results from ignoring any dim neighboring particles. Finally, in LD or HD mode, a counting error occurs when the inter-particle distance is below*d*_*HD/NR*_ (SNR).

**Figure 5.**
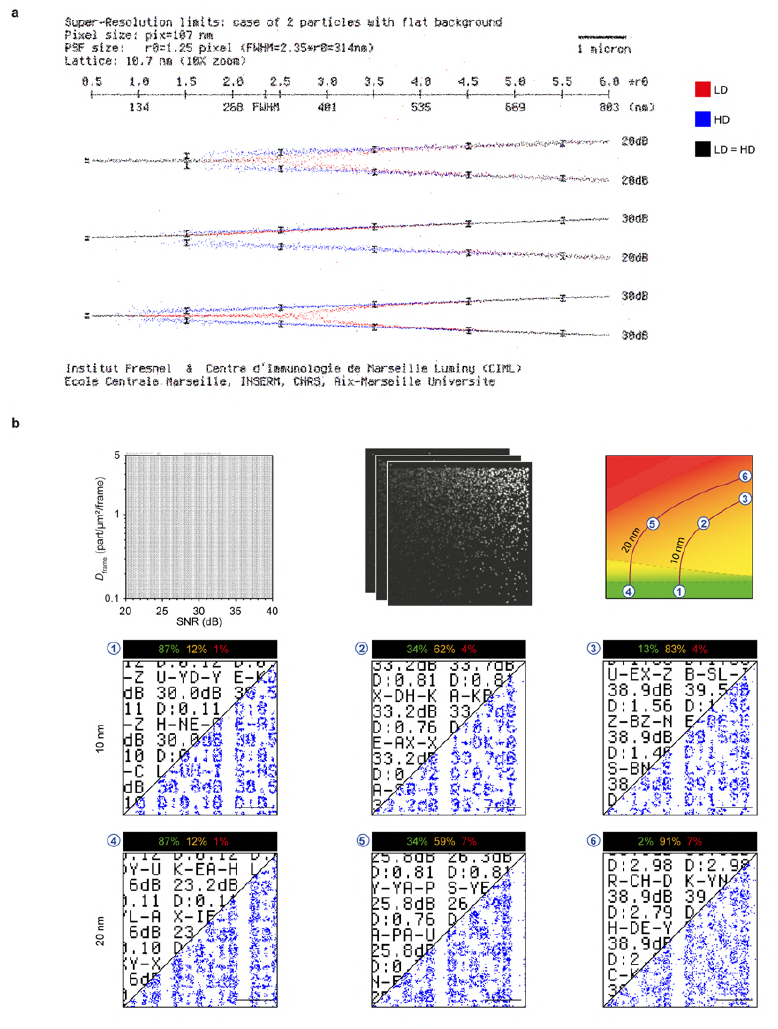
│ Evaluation of the performance of UNLOC in analyzing synthetic data. (**a**) The inter-particle distance chart mimics a two-particle canonical scenario at different SNR ratios. Binary renderings of the LD and HD modes of UNLOC are encoded in red and blue, respectively. Black pixels correspond to equivalent results for the LD and HD modes. As expected, the bias for particle localization appears at ≈ 5*r*_0_, the LD/HD transition limit and is higher for the LD mode, as shown by the dispersion around the lines. The HD/NR transition limit is clearly dependent on the SNR as shown for the different SNR ratio conditions. (**b**) The alphanumeric character chart reproduces the density-SNR space diagram (see Materials and Methods). Three ground truth areas and their corresponding binary rendering UNLOC results are determined along the 80% quantile isocurves at a localization precision of 10 and 20 nm. At a constant value for the localization precision, the bias is evidenced by a blurring effect, caused by the variation of LD/HD/NR subset partition along an isocurve. Scale bars: 500 nm.

The UNLOC analysis of the alphanumeric density-SNR chart leads to immediately visualize that the localization precision is preserved along an isocurve of the alphanumeric character chart (Fig. 5b). Compared to the standard ThunderSTORM HD algorithm (25), UNLOC provides more reliable estimates on such synthetic data concerning the localization precision as well as achievable resolution (47) (Figs. S12, S13). Moreover, UNLOC differs from super-resolution algorithms based on local density distribution reconstruction such as SRRF (7). As illustrated by the image reconstruction of experimental data for actin filaments in fixed cells, UNLOC efficiently challenges the background and density variability throughout the stack of images in a range of SNR of 18 to 40 dB with a localization precision ranging from 60 to 10 nm (Figs. 6, S12).

**Figure 6.**
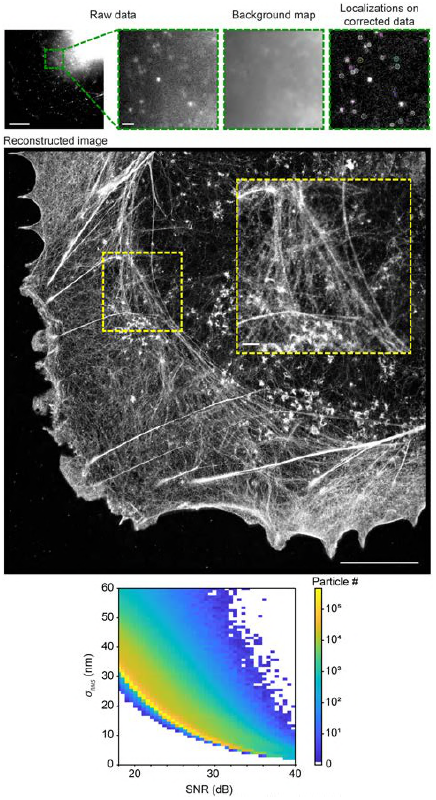
│ UNLOC analysis of dSTORM data. Alexa Fluor^™^ 647-phalloidin staining of actin filaments in COS-7 cells. Upper panels: the background map is estimated and subtracted from the raw data before the detection/enumeration step. The circle size depicts the localization precision. Middle panel: integrated Gaussian reconstructed image from ≈ 57,000 frames of 512 × 512 pixels each. Bars: 10 μm (insets, 1 μm). Lower panel: dot plot of the localization precision and SNR distributions corresponding to the reconstructed image.

Our theoretical study is intended for a homogeneous distribution for randomly distributed particles. However, experimental data deviate strongly from these ideal conditions. More generally, any biological study encounters possible high local densities, even at a low density per frame, and background variations, which fully justifies using an accurate HD mode of analysis. Such case is illustrated through the analysis of particles localized by UNLOC in LD versus HD mode for non-homogeneous distributed data (e.g., molecular clustering) (Fig. 7). The results of the clustering analysis by two different methods (31, 32) are recapitulated in Table S2. A proper quantification of clusters depends on the accuracy of the localization precision, which depends on local density, i.e., the density surrounding the particle in a given frame of the image stack. Of interest, UNLOC can also serve as an efficient localization tool for single particle tracking and the result further analyzed with the MTT algorithm (40).

**Figure 7.**
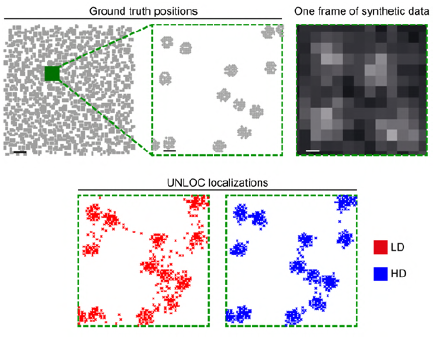
│ LD and HD modes of UNLOC analysis on non-homogeneous distribution. A synthetic stack of multiple frames mimics clustered ground truth positions with in green, a representative zoomed area of one frame (upper right). Lower panels: reconstructed image of the zoomed area analyzed in LD (red dots) or HD modes (blue dots) (see also Table S1). Bar: 1 μm (insets, 100 nm).

Finally, the classification of subgroups during the detection/estimation step does not modify the result but reduces the computation time by independently analyzing each individual subgroup with the appropriate fitting model. As order of magnitude, UNLOC analyzes in HD mode the density-SNR space diagram chart in ≈ 11 min (i.e., ≈ 20 frames/s or ≈ 3,200 particles/s). For an experimental biological data set of ≈ 57,000 frames of 512 ×512 pixels with non-homogeneous background and high variability of local density/frame (Fig. 6), UNLOC performs the task in HD mode within ≈ 200 min (i.e., ≈ 5 frames/s or ≈ 2,000 particles/s, for a total of ≈ 20 × 10^6^ detected particles).

## CONCLUSION

The strength of SMLM relates on a rigorous approach all along the experimental procedure, from the sample preparation to the data quantification. Here, we aim at assessing the accuracy and robustness of SMLM analysis by establishing a thorough theoretical study. We demonstrate that not only the SNR but also the local *D_frame_* strongly impact the real localization precision. More specifically, any neighboring particle strikingly biases the localization precision of a particle below an unexpected long inter-particle distance (≈ 600 nm). Also, even particles of weak SNR need to be detected since they contribute to ascertain unbiased localization of any neighboring particles. We identify inter-particle distance thresholds defining theoretically the working range of any localization-based algorithms leading to the definition of LD/HD/NR subsets, knowing the SNR-density values. In any case, non-resolvable scenario arises with increased local density/frame at an inter-particle distance *d*_*HD/NR*_ ≈ 170 nm, for two particles at 25 dB. Therefore, any NR scenario, which is by definition undeterminable with experimental data, impedes the localization precision, image representation, quantification and even the algorithm benchmarking performed with classical metrics. Moreover, we report that the knowledge of the particle intensity distribution does not improves the localization precision. Finally, our theoretical study clearly demonstrates that, compared to a MLE, an MMSE estimator is more robust and simplest to compute on high local density/frame data with complex background.

Therefore, we provide new tools that would help to report accurate quantitative SMLM observations. The density-SNR space diagram established for a realistic range of *D_frame_* and SNR enables standardized evaluation of the localization accuracy. Then, we develop UNLOC, a flexible and efficient unsupervised algorithm for high-density data. UNLOC is based on a heuristic insuring an optimal localization precision and counting of particles by considering both the SNR and *D_frame_* Finally, we also create synthetic data charts allowing blind benchmarking of any SMLM algorithms aiming specifically at providing a list of localizations of single molecules and not only a reconstructed image.

Altogether, this work provides an understanding of the key features usually underestimated or incorrectly taken into account in SMLM analyses.

## SUPPORTING MATERIAL

Fourteen figures, three tables, six notes and one software package accompany this paper.

**For referees, please use the temporary link below to download anonymously the Software package:**

https://filesender.renater.fr/?s=download&token=c679ccb6-b562-f543-0582-f0657de1b683

**This link is valid until February 15th, 2018.**

## AUTHORS’ CONTRIBUTIONS

N.B. and D.M. supervised the study and conceived the project. S.Ma., A.R., L.D., J.T. and N.B. developed the algorithms and performed the simulations; S.B., S.Ma. and S.Mo. designed the optical bench; R.F.,M.C.B. and Y.H. performed experimental observations; and all authors contributed to the interpretation of the data. N.B., S.Ma. and D.M. wrote the manuscript. The authors declare no competing financial interests.

## ACKNOWLEDGMENTS

We acknowledge Marc Allain, Sophie Brasselet, Hai-Tao He, Hervé Rigneault and Muriel Roche for their critical reading of the manuscript. We thank Christophe Leterrier for valuable discussions and Florian Levet for his advices on the SR-Tesseler analysis. This work was supported by institutional funding from the CNRS, Inserm, Aix-Marseille University and Centrale Marseille and program grants from the Research National Agency (ANR-10-BLAN-1214 to N.B. and D.M.), the French “Investissements d’Avenir” (ANR-11-IDEX-0001-02 A*MIDEX to N.B. and D.M., ANR-10-INBS-04 France BioImaging, ANR-11-LABX-0054 Labex INFORM and ANR-14-CE09-0008-02 to D.M.), the Fondation pour la Recherche Médicale (FRM-DEQ-20090515412 to D.M.) and the Institut National du Cancer (C15005AS to D.M.).

